# Relative adrenal insufficiency is a risk factor and an endotype of sepsis - A proof of concept study to support a precision medicine approach for glucocorticoid sepsis therapy

**DOI:** 10.1101/2020.04.16.043976

**Authors:** Chia-Hua Wu, Ling Guo, Qian Wang, Xiang Ye, Chieko Mineo, Philip W. Shaul, Xiang-An Li

**Affiliations:** The Department of Pharmacology and Nutritional Sciences, University of Kentucky College of Medicine, Lexington; Saha Cardiovascular Research Center, University of Kentucky College of Medicine, Lexington; Department of Pediatrics, University of Texas Southwestern Medical Center, Dallas; Department of Physiology, University of Kentucky College of Medicine, Lexington

**Keywords:** adrenal insufficiency, cecal ligation and puncture, corticosteroid therapy, inducible glucocorticoid, scavenger receptor BI, sepsis

## Abstract

**Rational:** 25-60% of septic patients experience relative adrenal insufficiency (RAI) and glucocorticoid (GC) is frequently used in septic patients. However, the efficacy of GC therapy and whether the GC therapy should be based on the status of RAI are highly controversial. Critical barriers include technical limitations in properly identifying RAI in septic patients and a lack of RAI animal model.

**Objectives:** We established a new RAI animal model to test our hypothesis that precision medicine approach should be used for GC sepsis therapy - only applying GC to a subgroup of septic mice with RAI.

**Methods:** We generated SF1CreSR-BI^fl/fl^ conditional knockout mice. The mice exhibited specific depletion of SR-BI expression in adrenal gland, resulting in a lack of production of inducible GC in response to ACTH stimulation or sepsis, but the mice had normal basal GC levels. Mice were treated with cecal ligation and puncture to develop sepsis. Mice were also supplemented with or without GC to study the effect of GC in sepsis therapy. Plasma and organs were collected for biochemical assays. BODIPY ^™^ FL-conjugated *Escherichia coli* was used for phagocytosis assay. Macrophages were used to study effects of GC on inflammatory responses.

**Measurements and Main Results:** Using SF1CreSR-BI^fl/fl^ mice as a RAI model, we found that mice with RAI were susceptible to CLP-induced sepsis compared to controls (6.7% survival in SF1CreSR-BI^fl/fl^ mice versus 86.4% in SR-BI^fl/fl^ mice; p=0.0001). Supplementation of hydrocortisone significantly improved survival in CLP-treated SF1CreSR-BI^fl/fl^ mice. Surprisingly, wild type mice receiving GC treatment exhibited significantly less survival compared to wild type mice without GC treatment. We further found that, in contrast to wild type mice which displayed a well-controlled systemic inflammatory response, the mice with RAI featured a persisted systemic response as shown by high levels of plasma inflammatory cytokines/chemokines 20 hours post CLP, and supplementation of GC kept the inflammatory response under control. In vitro analysis revealed that stress level of GC is required to suppress inflammatory response through modulating MAPK signaling in macrophages.

**Conclusions:** We demonstrate that RAI is a risk factor and an endotype for sepsis, and GC treatment benefits mice with RAI but harms mice without RAI. We further demonstrate that inducible GC functions to keep the systemic inflammatory response under control through modulating MAPK signaling, but mice with RAI lose such protection and supplementation of GC regains the protection. Our study provides a proof of concept to support the use of a precision medicine approach for sepsis therapy – selectively applying GC therapy for a subgroup of patients with RAI.

## Introduction

Sepsis affects 19 million people per year globally, with a mortality rate of ~ 30%^1–3^ Sepsis is caused by a dysregulated host response to infection. Upon infections, immune effector cells sense danger signals and generate inflammatory mediators to fight infections. However, over production of inflammatory mediators (hyper-inflammatory response) causes organ injury and under production of inflammatory mediators (hypo-inflammatory response) fails fighting infections.^4–8^ Thus, maintaining a balanced inflammatory response is critical for fighting infections as well as for avoiding organ injury.

Extensive efforts blocking one or another inflammatory mediator were not successful for septic patients, due to a number of potential limitations: **i)** there are a variety of inflammatory mediators; blocking one may not effectively control inflammation or maintain a balanced inflammatory response; **ii)** sepsis is complex and not all patients have over-inflammatory response. The anti-inflammatory therapy to patients without over-inflammation may cause immunosuppression. In light of the complexity of septic patients, emerging voices, including ours, have called for a precision medicine approach for sepsis therapy^9,10^. In support of this notion, a recent retrospective analysis showed that Anakinra (interleukin-1 receptor blocker), which showed no survival benefits for all septic patients in previous clinical trials,^11–13^ significantly improves 28-day survival in a subgroup of septic patients with features of macrophage activation syndrome.^14^ Therefore, identifying endotypes of sepsis is a key step for a successful precision medicine therapy for sepsis.

Adrenal insufficiency is commonly seen in septic patients ^15–17^, especially relative adrenal insufficiency (RAI) ^18,19^. RAI is considered as the insufficient production of inducible cortisol (iGC) for the current physiologic stress, no matter what the level of plasma cortisol is ^20–22^ The impaired adrenal function could be a potential risk factor for mortality in septic patients. Septic patients who failed to produce iGC in response to ACTH stimulation were susceptible to sepsis-induced death ^18,23,24^, which indicates the critical roles of iGC in sepsis. Since the critical roles of adrenal insufficiency in sepsis, numerous clinical trials for corticosteroid therapy have been conducted. However, the evidence of mortality benefit of corticosteroid therapy in sepsis is still inconclusive ^25,26^. An early study by Annane et al. ^18^ showed that low-dose of corticosteroid treatment significantly reduced the mortality of septic shock patients with relative adrenal insufficiency^18^. However, a later CORTICUS trial failed to validate these findings^17^ in a less severe group of septic patients with RAI. HYPRESS trial tested the efficacy of hydrocortisone in patients with sepsis without shock and found that the use of hydrocortisone did not reduce the risk of septic shock within 14 days or improve survival^27^. The ADRENAL trail involving 3800 septic shock patients (without distinguishing the status of RAI) did not show improvement in survival^28,29^. The APROCCHESS trial reported that the use of hydrocortisone plus fludrocortisone improved the 90 day survival in patients with septic shock^30^. Nevertheless, the debate is going on. Surviving Sepsis Campion Guidelines suggest using hydrocortisone as a treatment of adult septic shock patients who are refractory to vasopressor therapy (weak recommendation, low quality of evidence) ^31^. Despite the controversial clinical outcomes of corticosteroid therapy and weak recommendation, low-dose corticosteroid is often used for septic shock patients (50.4%) ^32^ These data show that (1) the lack of proper method of identifying RAI is a barrier of corticosteroid therapy used in septic patients, and (2) a demand for corticosteroid in septic patients increases even though a lack of clear evidence indicating the mortality benefit, which points out an urgent need to determine the efficacy of corticosteroid therapy.

We hypothesize that RAI is an endotype of sepsis and GC therapy benefits patients with RAI. To address this hypothesis, we generated a specific mouse model having RAI, called SF1creSR-BI^fl/fl^ mice (adrenocortical SR-BI knockout mice). Recent studies of our laboratory and others reported that scavenger receptor class B type I (SR-BI), a high-density lipoprotein (HDL) receptor facilitating the uptake of cholesterol from HDL into target tissues ^33,34^, is critical for the protection against sepsis ^35–38^ and also essential for GC production in adrenal glands ^35,39,40^. In the present study, we demonstrated that iGC modulates the inflammatory responses in sepsis. The physiological level of GC is not adequate for the protection against sepsis. GC therapy is beneficial for the survival of mice with adrenal insufficiency but not wild-type mice. Collectively, our study suggests that GC therapy could be a potential sepsis treatment for patients with relative adrenal insufficiency.

## Material and method

### Mice

SF1cre mice in FVB/NJ background were from the Jackson Laboratory (#012462). SR-BI ^fl/fl^ mice in 8x C57BL/6J background were from Dr. Chieko Mineo (University of Texas Southwestern Medical Center). SF1cre mice were backcrossed for 4 generations against C57BL/6J mice and subsequently interbred with SR-BI ^fl/fl^ mice to yield SF1creSR-BI ^fl/fl^ mice (FVB/NJ x C57BL/6J). The animals were bred at the University of Kentucky’s animal facility. The animals were fed a standard laboratory diet and kept with a 10/14 h light/dark cycle and littermates were used at 10-14-week old. Animal care and experiments were approved by the Institutional Animal Care and Use Committee of the University of Kentucky.

### Cecal ligation and puncture (CLP)

CLP was performed following the protocol described previously ^37^ Briefly, 10- to 14-week-old mice were anesthetized by inhalation of 2–5% isoflurane in 100% oxygen. A midline incision (~1.0 cm) was made on the abdomen. The cecum was ligated and punctured twice with a 25-gauge needle, and gently compressed to extrude a small amount of cecal material. The cecum was returned to the abdomen, and the incision was closed with 6–0 Ethilon suture material. The mice were subsequently resuscitated with 1 ml phosphate-buffered saline via i.p. Survival was monitored for a 7-day period.

### Assay for SR-BI Expression

Tissues were homogenized in lysis buffer containing 1% proteinase inhibitor mixture (Sigma), and subjected to Western blot analysis against SR-BI as described previously ^41^. The expression of SR-BI was quantified with Fuji LAS-4000 and normalized to actin expression.

### Biochemical assays

Mice were sacrificed by CO_2_ inhalation and plasma was obtained by cardiac puncture and stored at −80 °C. The plasma corticosterone was quantified with a kit from ENZO Life Sciences (cat# ADI-900-097). Cytokines (IL-6 and TNF-α) were quantified with corresponding ELISA kits from eBioscience (cat# 88-7064 and cat# 88-7324, respectively). Plasma cytokines were also analyzed by Mouse Cytokine Array / Chemokine Array 31-Plex (MD31) which was provided by Eve Technologies. The serum alanine aminotransferase (ALT) levels were quantified with a kit from POINTE SCIENTIFIC, Inc (cat# A7526-01-881) to determine the degree of liver damage. The blood urea nitrogen (BUN) concentrations were measured with a kit from the QuantiChrom to determine the degree of kidney injury (cat# DIUR-100).

### Assay for phagocytosis of bacteria

BODIPY ™ FL-conjugated *Escherichia coli* BioParticles (*E. coli*, K-12 strain, Life technologies) were suspended in sterile PBS to a final concentration of 1 ×10^9^ CFU/ml. The BODIPY-*E. coli* were intraperitoneally injected at 1 ×10^8^/mouse at 17 h post CLP. One hour later, the peritoneal fluid, blood and spleen were harvested for FCAS analysis.

### Fluorescence-activated cell sorting (FACS) analysis

FACS analyses were performed as we described previously ^42^. 2×10^5^ cells were blocked with anti-mouse CD16/CD32 (Fc) and subsequently incubated with fluorochrome-conjugated antibodies from BD or Biolegend, including PE-, APC-, APC-cy7, PerCP-Cy5.5, or PE-Cy7-conjugated anti-CD3 (17A2), CD11b (M1/70), CD11c (HL3), CD45 (30F11), F4/80 (BM8), and Ly6g (1A8) for 30 min at 4 °C. The stained cells were then washed and resuspended in the FACS buffer and analyzed with LSRII Flow Cytometer (BD). The FACS data were analyzed with FlowJo software (Treestar).

### Supplementation of corticosterone in mice

Hydrocortisone (water-soluble, Sigma-Aldrich, cat# H0396) was dissolved in PBS and i.p. injected at 75 μg/mouse immediately following CLP. Survival was monitored for a 7-day period.

### Cell Culture

J774 macrophage cell line (ATCC) were maintained in DMEM containing 10% FBS (Premium Feta Bovine Serum, Atlanta), 2% HEPES (pH 7.4), 1% glutamine and 1% Pen/Strep and cultured at 37°C in a 5% CO_2_ in air atmosphere. Bone marrow-derived macrophages were cultured as described previously ^43^. Bone marrow-derived macrophages (BMDM) were isolated from SR-BI+/+ (129×C57BL/6) mice and cultured in RPMI 1640 containing 10% FBS (Premium Feta Bovine Serum, Atlanta), 1% glutamine,1% Pen/Strep and 15% L929 medium for 7 days. The medium was changed once every three days.

### Quantification of cytokine generation in LPS-stimulated macrophages

For J774 cells, cells (4×10^4^ cell/well; 96 well plate) were cultured for two days and treated with 0, 125, 250, 500, 1000, 2000 and 4000 nM hydrocortisone (lipid-soluble, Sigma-Aldrich) in the presence of 2 ng/ml LPS (*Escherichia coli* serotype K12, InvivoGen) for 18h. The supernatants were harvested and analyzed for IL-6 and TNF-α concentration. For BMDM, cells (5×10^5^ cell/well; 6 well plate) were cultured for 7 days and treated with or without 0.4ng/ml IFN-γ and with 0, 32.25, 100, 500 and 2000 nM hydrocortisone in the presence of 2 ng/ml LPS for 18h. The supernatants were harvested for cytokine analysis and cells were harvested for qRT-PCR.

### Analysis of gene expression by quantitative reverse transcription PCR

Total RNA was isolated from cells using RNeasy Mini Kit (QIAGEN), following the manufacturer’s instructions. cDNA was synthesized from 100 ng of total RNA, using iScript™ Reverse Transcription Supermix (Bio-Rad). PCR amplification of the cDNA was performed by qPCR using iQ™ SYBR®

Green Supermix (Bio-Rad). The thermocycling protocol consisted of 3 min at 95 °C, 40 cycles of 15 sec at 95 °C, and 45 sec at 58 °C, and finished with a melting curve ranging from 60 to 95 C to allow distinction of specific products. qPCR were performed with primers of MKP-1 (5’- GTTGTTGGATTGTCGCTCCTT and 5’-TTGGGCACGATATGCTCCAG) and U36B4 (5’- CGTCCTCGTTGGAGTGCA and 5’-CGGTGCGTCAGGGATTG). U36B4 RNA was amplified as an internal control.

### Statistical Analysis

The survival assay was analyzed by Log-Rank χ^2^ test. Significance in experiments comparing two groups was determined by 2-tailed Student’s t test. Significance in experiments comparing more than two groups was evaluated by One Way ANOVA, followed by post hoc analysis using Tukey’s test. Means were considered different at *p*< 0.05.

## Result

### Mice with adrenal insufficiency were susceptible to CLP-induced septic death

It was reported that adrenal SR-BI is required for the production of inducible glucocorticoid (iGC) in sepsis in a relative adrenal insufficiency mouse model generated by adrenal transplantation ^38^. Adrenal transplantation has few limitations such as technique issue and the cut-off of the neuron and chromaffin cells communication. Therefore, we generated SF1creSR-BI^fl/fl^ mice (adrenocortical SR-BI knockout mice) as another relative adrenal insufficiency animal model by breeding SF1-cre mice with SR-BI^fl/fl^ mice (Fig 1A). Firstly, we identified whether SF1creSR-BI^fl/fl^ mice is adrenal insufficiency. Western blot analysis of adrenal extracts from SF1creSR-BI^fl/fl^ mice showed undetectable levels of SR-BI protein; however, the level of SR-BI protein did not change in liver and adrenal extract from and SR-BI^fl/fl^ mice as well as liver extract from SF1creSR-BI^fl/fl^ mice, showing that SR-BI was specifically knocked out in adrenal glands and *loxP* sites insertion did not affect SR-BI expression globally (Fig 1B). At the basal physiologic condition, plasma corticosterone concentration did not show significant difference between SR-BI^fl/fl^ mice and SF1creSR-BI^fl/fl^ mice. After ACTH stimulation, SF1creSR-BI^fl/fl^ mice failed to generate inducible corticosterone, compared with SR-BI^fl/fl^ mice (Fig 1C). These data indicate that SF1creSR-BI^fl/fl^ mouse is a useful model of relative adrenal insufficiency.

**Fig1.**
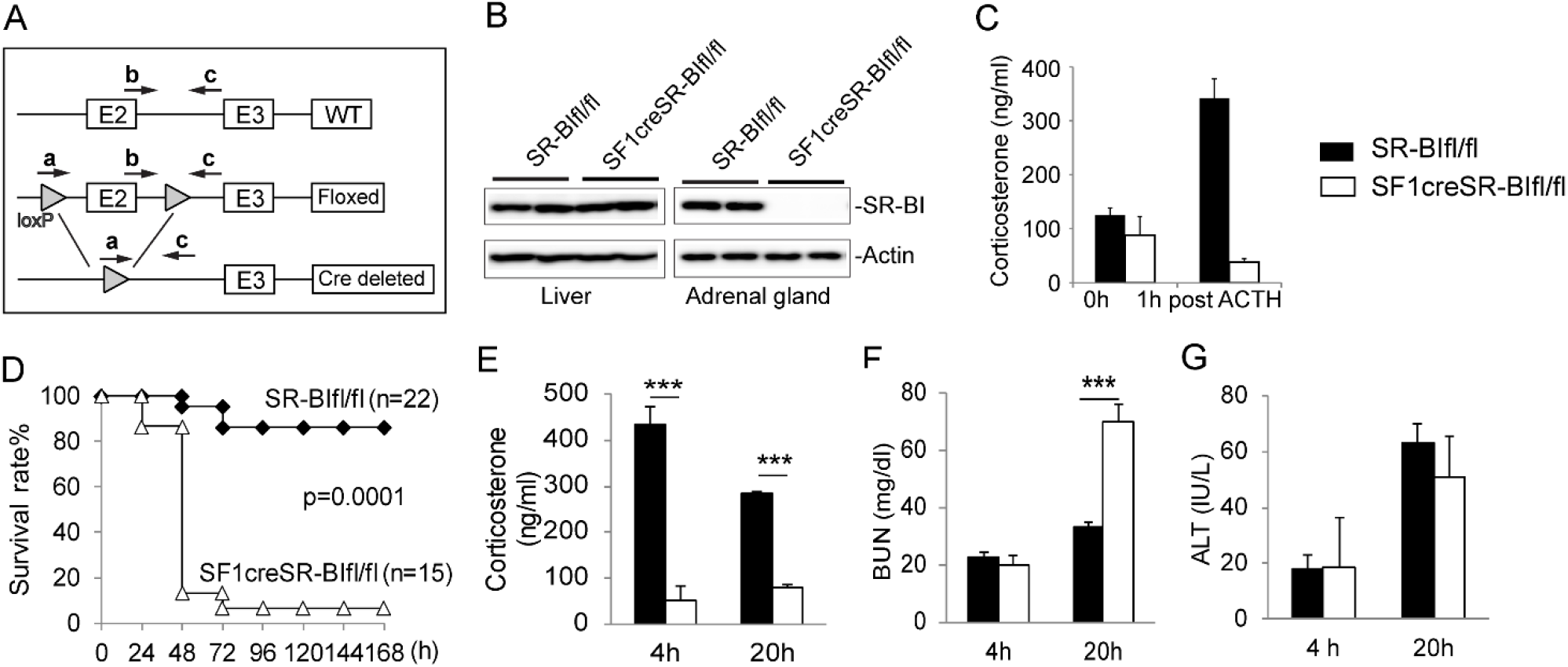
Adrenal SR-BI knockout mice which were corticosteroid insufficiency were susceptible to CLP-induced septic death. (A) A schematic of the Cre-lox system used to generate SF1creSR-BI^fl/fl^ mice. a, b and c represent as primers. E2 and E3 represent as exon 2 and exon 3, respectively. (B) Western blot analysis of SR-BI expression in the liver and adrenal gland in SR-BI^fl/fl^ mice and SF1creSR-BI^fl/fl^ mice. (C) Plasma corticosteroid levels 1h post adrenocorticotropic hormone (ACTH) stimulation in SR-BI^fl/fl^ mice and SF1creSR-BI^fl/fl^ mice. (D) Survival analysis. SR-BI^fl/fl^ mice and SF1creSR-BI^fl/fl^ mice were treated with CLP (25G, full ligation) and monitored for 7 days for survival analysis (n=15-22). Data are expressed as the percentage of mice surviving at the indicated time points and analyzed by the log-rank χ2 test. SR-BI^fl/fl^ mice and SF1creSR-BI^fl/fl^ mice were treated with CLP (25G, full ligation) for 4 and 20h. Plasma was harvested and analyzed for (E) plasma corticosterone levels, (F) blood urea nitrogen (BUN) levels as a marker for kidney injury and (G) plasma alanine aminotransferase (ALT) levels as a marker for liver injury. n = 5-8 for (E-G). Data represent means ± sem. ***p<0.001 denote significant differences compared to SR-BI^fl/fl^ mice by t test.

Mice with adrenal insufficiency are susceptible to CLP-induced septic death and kidney injury. As shown in Figure 1D, SF1creSR-BI^fl/fl^ mice showed significantly higher mortality compared with SR-BI^fl/fl^ mice (*p*<0.0001). Similar to data from ACTH stimulation, serum corticosterone level significantly increased in SR-BI^fl/fl^ mice at 4 and 20 h after CLP treatment. On the contrary, SF1creSR-BI^fl/fl^ mice failed to increase serum corticosterone concentration (Fig 1E). Organ injury is a hallmark of sepsis. To understand whether organ injury is associated with CLP-induced sepsis death, markers of liver and kidney injuries were assayed. Blood urea nitrogen concentration was about 2 times higher in SF1creSR-BI^fl/fl^ mice than SR-BI^fl/fl^ mice 20 h after CLP treatment (Fig 1F). However, there was no significantly difference in serum alanine aminotransferase level between SF1creSR-BI^fl/fl^ mice and SR-BI^fl/fl^ mice (Fig 1G). These data show that the deficiency of GC leads to kidney injury and death in CLP-induced sepsis model.

### Mice with adrenal insufficiency had uncontrolled cytokine release and impaired phagocytotic ability in sepsis

Since glucocorticoid modulates a broad spectrum of inflammation-related genes and signaling pathways, it is an important part of resolution of inflammation ^44^. To investigate whether the immune responses are affected in our adrenal insufficiency mouse model in sepsis, we measured 31 plasma cytokines (Mouse Cytokine Array / Chemokine Array 31-Plex; MD31) and analyzed the phagocytic capability of phagocytes in CLP-treated mice. Interestingly, all analyzed cytokines except for IL-4, IL-7 and IL-12(p70) were higher in SF1creSR-BI^fl/fl^ mice than SR-BI^fl/fl^ mice at 20 h after CLP treatment (Supplemental Table S1 and Fig 2). Especially Eotaxin, GM-SCF, IL-1B, IL-2, IL-5, IL-6, IL-12(p40), IL-13, KC, MIP-1alpha, TNF-alpha and VEGF were significantly high in SF1creSR-BI^fl/fl^ mice at 20 h after CLP treatment compared with SR-BI^fl/fl^ mice (Fig 2). In summary, it shows that the deficiency of iGC causes systemic hyper-inflammatory responses in sepsis.

**Fig2.**
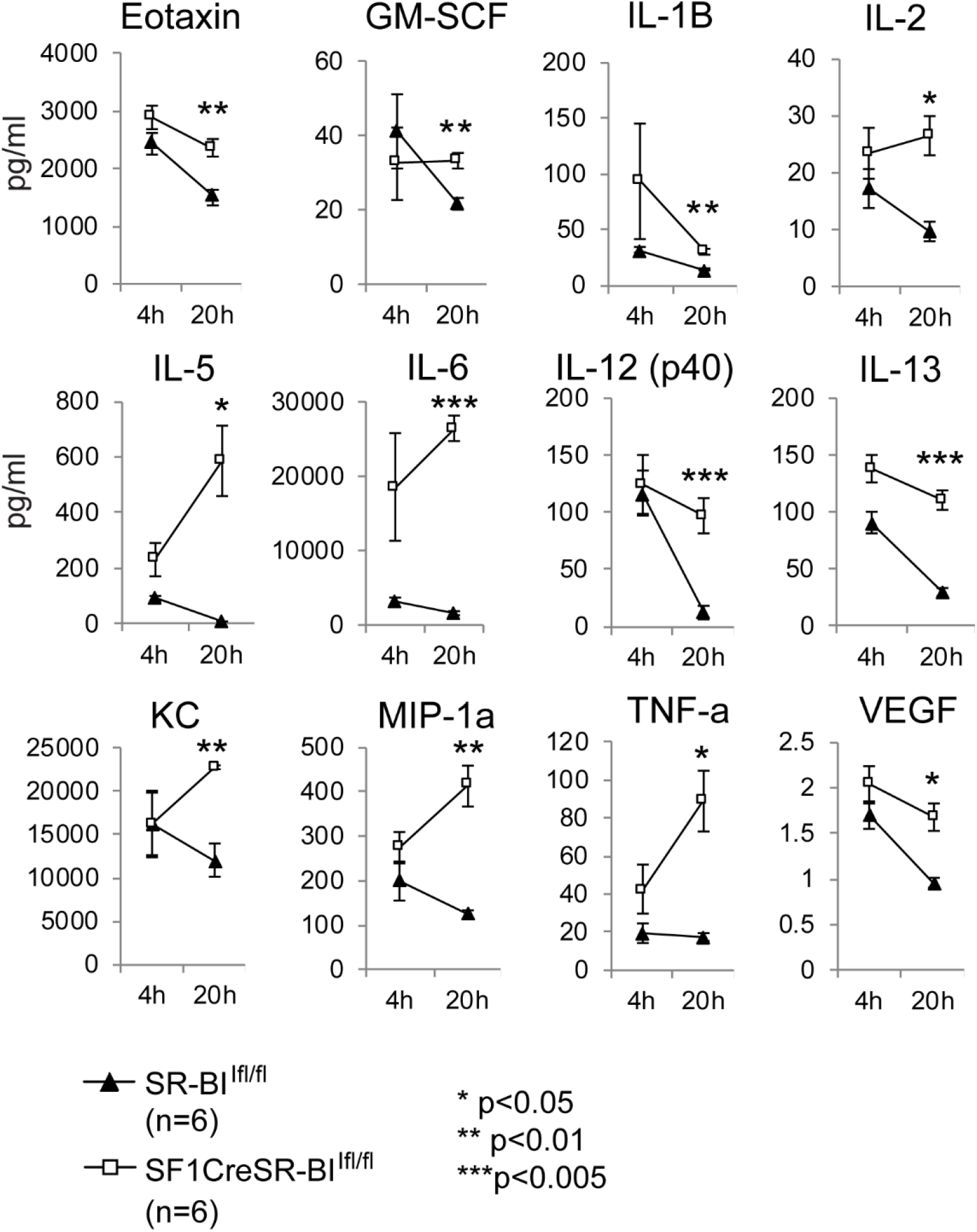
Mice with adrenal insufficiency had hyper-inflammatory response in sepsis. SR-BI^fl/fl^ mice and SF1creSR-BI^fl/fl^ mice were treated with CLP (25G, full ligation) for 4 and 20h. Plasma was harvested and analyzed for cytokines levels. Data represent means ± sem. * p < 0.05, ** p < 0.01 and *** p < 0.005 denote significant differences compared to SR-BI^fl/fl^ mice by t test.

We utilized BODIPY ™ FL-conjugated *Escherichia coli* BioParticles to test the phagocytosis in sepsis. As shown in Fig 3, in peritoneal fluids, compared with the control mice, SF1creSR-BI^fl/fl^ mice had a 58% lower percentage of *E. coli+* CD11b+ cells (CD45+ CD11 b+; 54.3±4.57% vs 22.7±12.9%), a 86% lower percentage of *E. coli+* macrophages (CD45+ CD11b+ F4/80+; 69.5±3.94% vs 9.78±7.10%) and a 80% lower percentage of *E. coli+* neutrophils (CD45+ CD11b+ Ly6g+; 33.82±4.96% vs 6.8±6.44%). The phagocytic capability of phagocytes was evaluated by the mean fluorescence intensity (MFI) of *E. coli* in the phagocytes. In peritoneal fluids, compared with the control mice, SF1creSR-BI^fl/fl^ mice had a 60% lower percentage of MFI of *E. coli* in CD11b+ cells (17.05±1.86 x 10^3^ vs 6.87±3.27 x 10^3^, p=0.036), a 73% lower percentage of MFI of *E. coli* in macrophages (19.37±1.51 x 10^3^ vs 5.22±0.9 x 10^3^, p=0.0027), and a 56% lower percentage of MFI of *E. coli* in neutrophils (17.26±2.72 x 10^3^ vs 7.56±3.10 x 10^3^, p=0.1)(Fig 3). However, the phagocytic capability of neutrophils in blood and spleen did not significantly change (Supplemental Fig S1). In summary, these data show that the deficiency of iGC impairs the phagocytic ability of phagocytes in sepsis.

**Fig3.**
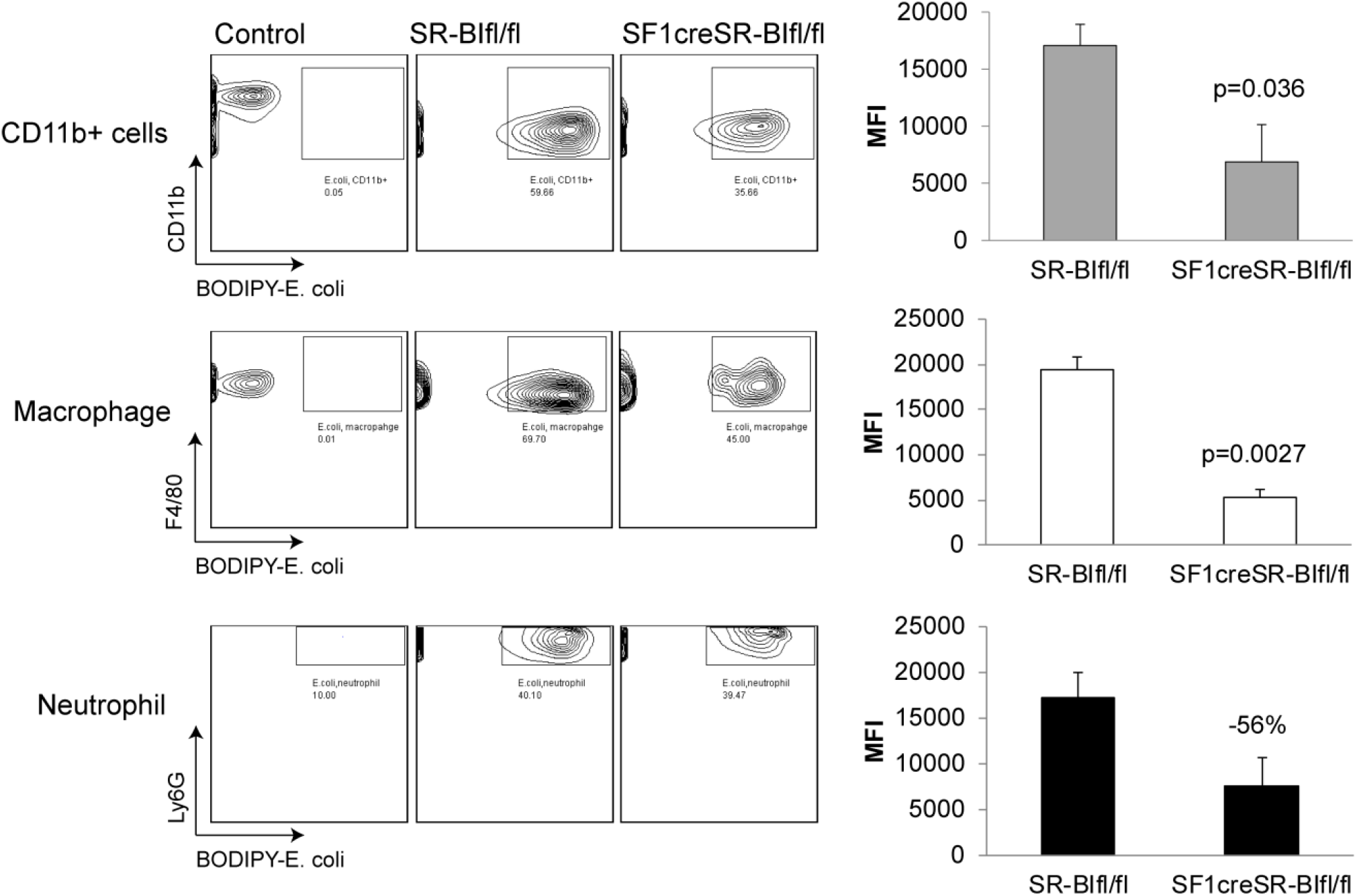
The deficiecny of iGC impaired the phagocytic ability of phagocytes in sepsis. SR-BI^fl/fl^ mice and SF1creSR-BI^fl/fl^ mice were treated with CLP (25G, full ligation) for 17h and then injected with 10^9^ CFU/ml BODIPY-conjugated *E. coli* via i.p. After 1 hour, the mean fluorescence intensity (MFI) of *E. coli* in phagocytes in peritoneal fluids was analyzed with flow cytometry. n = 2-4. Data represent means ± sem. Control represents mice without CLP and *E. coli* treatment. * p < 0.05 denotes significant differences compared to SR-BI^fl/fl^ mice by t test.

### GC supplementation benefits septic mice with adrenal insufficiency but harms septic mice without adrenal insufficiency

Since whether targeting adrenal insufficiency in septic patients for sepsis therapy is still controversial, we used SF1creSR-BI^fl/fl^ mice as a specific adrenal insufficiency animal model to investigate the effect of GC supplementation on survival of septic mice with adrenal insufficiency. Hydrocortisone is the pharmaceutical term for cortisol used in human medicine. As shown in Fig. 4A, 75 μg/mouse hydrocortisone treatment significantly decreased the survival rate of SR-BI^fl/fl^ mice. On the contrary, 75 μg/mouse hydrocortisone treatment significantly increased the survival rate of SF1creSR-BI^fl/fl^ mice (Fig. 4B). These data show that GC supplementation is beneficial for septic mice with adrenal insufficiency but not septic mice with normal adrenal function.

Interestingly, at 3 days post CLP, hydrocortisone treatment increased the survival of SF1creSR-BI^fl/fl^ mice (survival rate: non GC treatment group= 12.5%, GC treatment group= 76.9%); at 4 days post CLP, hydrocortisone also increased the survival of SF1creSR-BI^fl/fl^ mice (survival rate: non GC treatment group= 0%, GC treatment group= 61.5%). However, the survival rate of CLP-treated SF1creSR-BI^fl/fl^ mice injected with hydrocortisone decreased gradually, showing that early single dose of GC administration improved the survival of septic mice with adrenal insufficiency but the efficacy of single dose of GC reduced with the time course.

**Fig4.**
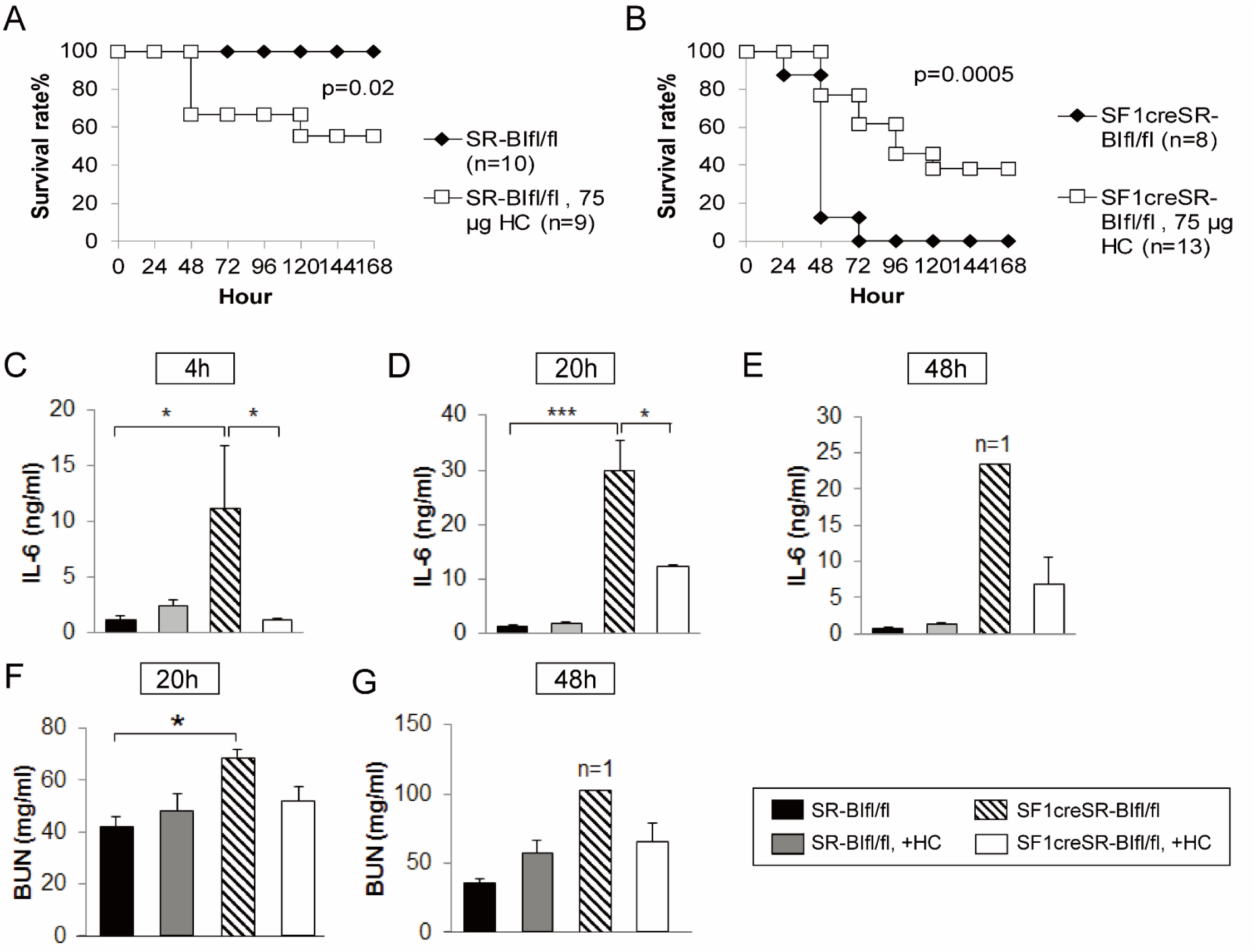
GC supplementation increased the survival rate of mice with adrenal insufficiency, but not mice without adrenal insufficiency. (A) SR-BI^fl/fl^ mice and (B) SF1creSR-BI^fl/fl^ mice were injected with or without 75μg/mouse hydrocortisone via i.p. after CLP (25G, full ligation) treatment. Mice were monitored for 7 days for the survival rate. Data are expressed as the percentage of mice surviving at the indicated time points and analyzed by the log-rank χ2 test. Blood were harvested from tail vein at 4, 20, 48 and 72h after treatment for the analysis of IL-6 concentration (C-E) and blood urea nitrogen (BUN) levels (F-G). n=5-9. Data are presented as the means ± sem. (C-G) * p < 0.05 and ***p<0.001 denote significant differences by one way ANOVA.

IL-6 is correlated with mortality and severity scores in septic patients ^45^. Also, acute kidney injury enhances risk of multiple organ failures and mortality ^46^. Therefore, to understand why GC therapy benefits septic mice with adrenal insufficiency, we analyzed the plasma level of IL-6 and BUN in these CLP-treated mice. As shown in Fig. 4C, 4D and 4E, plasma IL-6 concentration was significantly higher in SF1creSR-BI^fl/fl^ mice than SR-BI^fl/fl^ mice at 4, 20 and 48 h post CLP treatment. GC treatment significantly reduced plasma IL-6 concentration in SF1creSR-BI^fl/fl^ mice compared with non-GC-treated SF1creSR-BI^fl/fl^ mice at 4, 20 and 48 h post CLP treatment. In the meanwhile, SR-BI^fl/fl^ mice with GC supplementation displayed higher plasma level of IL-6 compared with non-GC-treated SR-BI^fl/fl^ mice until 72h post CLP. These data suggest that GC supplementation significantly reduced IL-6 production in mice with adrenal insufficiency during sepsis. Although there was no significant difference in IL-6 level between two groups of SR-BI^fl/fl^ mice, GC supplementation increased IL-6 level in SR-BI^fl/fl^ mice in sepsis.

It showed that the deficiency of iGC caused the CLP-induced kidney injury but not liver injury at 20h post CLP treatment in Fig. 1F and 1G. Thus, we also tested whether GC supplementation enabled to ameliorate kidney injury in sepsis. As shown in Fig. 4F, SF1creSR-BI^fl/fl^ mice had significantly high plasma blood urea nitrogen (BUN) level at 20h after CLP compared with SR-BI^fl/fl^ mice (*p*<0.05). GC treatment decreased 24% of BUN level in SF1creSR-BI^fl/fl^ mice at 20h after CLP treatment. Only one SF1creSR-BI^fl/fl^ mice survived at 48h post CLP treatment (Fig. 4G). Compared with this SF1creSR-BI^fl/fl^ mouse, SR-BI^fl/fl^ mice had 190% lower percentage of BUN and GC-treated SF1creSR-BI^fl/fl^ mice had 36% lower percentage of BUN (Fig. 4G). These data show that the deficiency of iGC results in severe kidney injury and GC supplementation improves this injury during sepsis. In addition, GC treatment increased 36% of BUN in SR-BI^fl/fl^ mice at 48h after CLP treatment (Fig. 4G), showing that GC treatment aggravates septic mice with normal adrenal function.

### Stress level of GC (iGC) is required for the effective suppression of inflammatory cytokine production in the presence of IFN-γ

Macrophages are critical immune cells involved in inflammatory cytokines production against infection in sepsis ^47^. Therefore, to investigate the relationship of GC concentration with the inhibitory effects on pro-inflammatory cytokines production in sepsis, we utilized macrophage cell lines and bone marrow-derived macrophages (BMDMs) to study the roles of GC level in inflammation in sepsis. As shown in Fig. 5A and 5B, hydrocortisone significantly reduced IL-6 and TNF-α production in LPS-treated J774 cells and BMDMs in a dose-dependent manner.

**Fig5.**
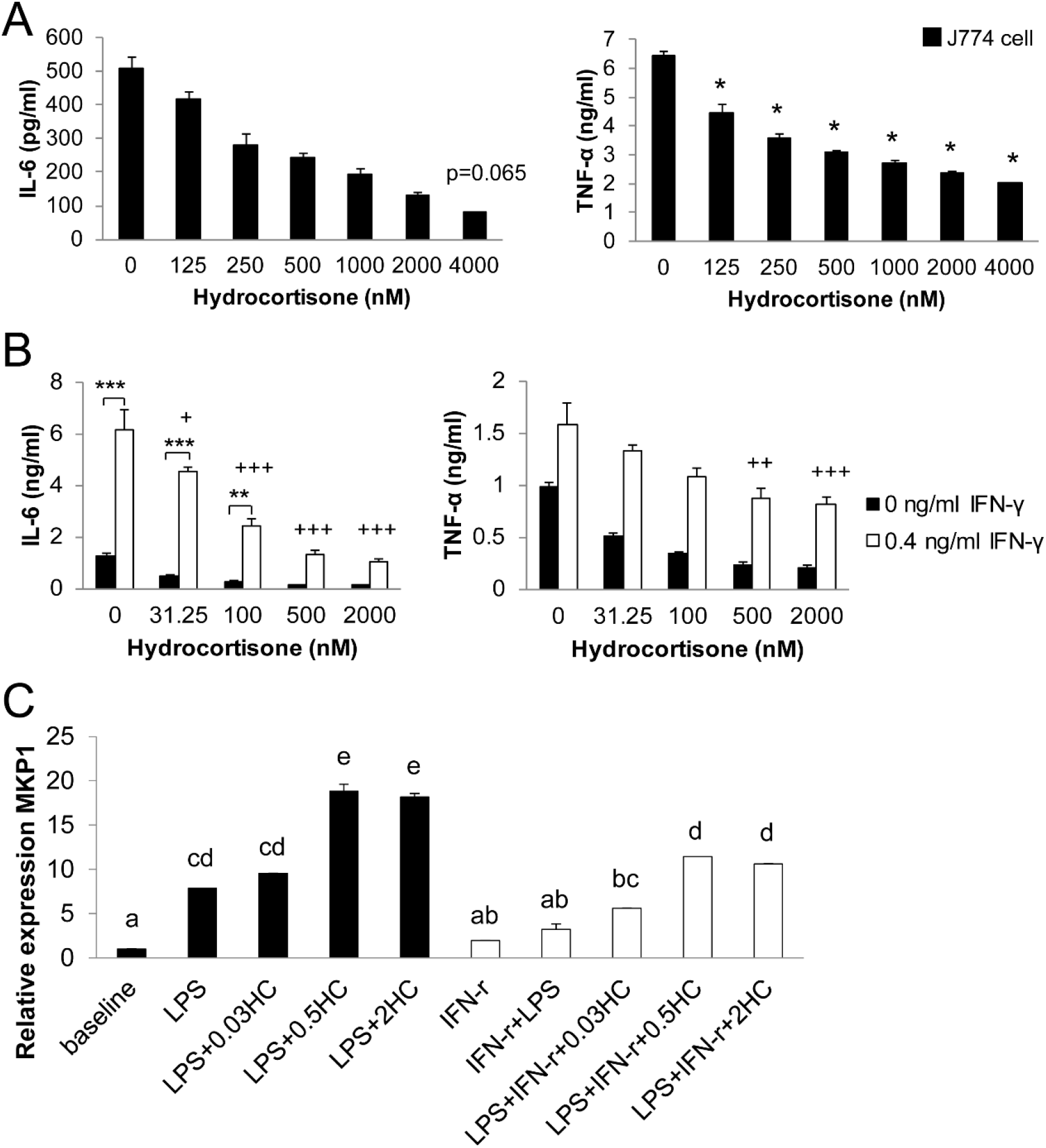
Stress level of GC is required for effective suppression of inflammatory cytokine production. (A) J774 cells were treated with 0, 125, 250, 500, 1000, 2000 and 4000 nM hydrocortisone in the presence of 2 ng/ml LPS for 18h. The supernatants were harvested and analyzed for IL-6 and TNF-α concentration. Data are presented as the means ± sem. * p < 0.05 denotes significant differences compared to non-hydrocortisone treatment by one way ANOVA. Bone marrow-derived macrophages (BMDM) were isolated from SR-BI+/+ mice in 129×B6 background. After 7 days cultured with complete DMEM-10 with 15% L929 medium, BMDMs were treated with or without 0.4ng/ml IFN-γ and with 0, 31.25, 100, 500 and 2000 nM hydrocortisone in the presence of 2 ng/ml LPS for 18h. The supernatants were harvested for cytokine analysis (B) and cells were harvested for qRT-PCR (C). Results were normalized to mouse U36b4 RNA expression. Data are presented as the means ± sem. (B)* p < 0.05, ** p < 0.01 and ***p<0.001 denote significant differences compared to non-IFN-γ treatment by one way ANOVA. + p < 0.05, ++ p < 0.01 and +++p<0.001 denote significant differences compared to non-HC treatment in IFN-γ treatment group by one way ANOVA. (C) a, b, and c denote significant differences by one way ANOVA with Tukey HSD test.

The higher concentration of GC is stimulated and required for higher stress condition in sepsis (Fig. 1E and 2). IFN-γ, secreted by Th1 T lymphocytes, is one of main macrophage activators ^48^. IFN-γ-priming increases the production of cytokines in LPS-treated macrophage. We utilized BMDMs treated with or without IFN-γ, LPS and hydrocortisone to investigate whether the higher level of GC is required for the inhibition of cytokine production in IFN-γ-stimulated macrophages. As shown in Fig. 5B, in IFN-γ co-stimulation, IL-6 and TNF-α production were significantly enhanced in LPS-treated BMDMs. Hydrocortisone significantly decreased IL-6 and TNF-α production in either LPS-treated BMDMs or LPS and IFN-γ co-stimulated BMDMs in a dose-dependent manner. However, in the IFN-γ co-stimulation, higher hydrocortisone is required for the inhibition of cytokine production in LPS-treated BMDMs compared with non-IFN-γ co-stimulated group.

Glucocorticoids induce several anti-inflammatory genes such as MAP kinase phosphatase-1 (MKP-1), which plays an important role in the feedback control of MAP kinase signaling ^49,50^. We next investigated the effect of GC on the expression of MKP-1 in the presence of IFN-γ. As shown in Fig. 5C, 0.03, 0.5 and 2 μM hydrocortisone increased the mRNA expression of MKP-1 in LPS-treated cells. Interestingly, IFN-γ only as well as IFN-γ and LPS co-stimulation did not significantly change the mRNA expression of MKP-1 compared with non-treated cells. 0.03 μM HC+LPS+IFN-γ group and 0.5 μM HC+LPS+IFN-γ group had 1.7 time and 3.5 times fold higher mRNA expression of MKP-1 compared with LPS+IFN-γ group. Compared LPS-treated groups with LPS+ IFN-γ groups, the synergistic effect of IFN-γ and LPS compromises the positive effects of GC on mRNA expression of MKP-1. These results show that stress level of GC is required for the effective suppression of pro-inflammatory cytokine production in LPS-stimulated macrophages in the presence of IFN-γ.

## Discussion

This study demonstrates that iGC is a critical survival factor of mice in sepsis and the supplementation of corticosteroid benefits the survival of septic mice with adrenal insufficiency. iGC is involved in the modulation of systemic inflammatory responses. iGC is required for the suppression of pro-inflammatory responses and amelioration of organ injury.

### Adrenal insufficiency worsens the outcomes of sepsis

Several studies showed that adrenal insufficiency increases the risk of death of septic patients ^17–19^,^51–53^. The variation of cortisol level was seen in sepsis. High basal plasma cortisol level and weak cortisol response to corticotropin stimulation (basal plasma cortisol>34μg/dl and Δmax≤9μg/dl) was positively associated with high 28-day mortality in septic patients ^19^. Another data showed that basal plasma cortisol>20μg/dl and Δmax≤9μg/dl resulted in significantly high mortality (82%) in septic patients^53^. In animal studies, mice with adrenal insufficiency were very susceptible to sepsis-induced death as well ^35,37,38,54^. These suggest that adrenal functions were strongly correlated with sepsis-related mortality. In this study, we successfully generated a relative adrenal insufficiency animal model (SF1creSR-BI^fl/fl^ mice). As expected, SF1creSR-BI^fl/fl^ mice which failed to produce iGC were susceptible to CLP-induced sepsis death and displayed aberrant inflammatory responses and kidney injury. Interestingly, among 32 cytokines we measured, most pro-inflammatory cytokines markedly increased compared with control mice (Fig.2), showing the critical roles of iGC in the modulation of inflammatory responses in sepsis.

Phagocytosis is part of critical defense of microbial infection in sepsis. The impaired phagocytosis aggravates sepsis^55^. It has been reported that GC enhances the phagocytic ability of phagocytes. The insufficiency of iGC impaired the phagocytic ability of monocytes, macrophages and neutrophils in blood in sepsis ^38^. In this study, we assessed the phagocytosis of *E. coli* in peritoneal cavity in addition to spleen and blood. SF1creSR-BI^fl/fl^ mice had significantly lower phagocytic ability in phagocytes in peritoneal cavity, although there was no significant difference in phagocytic ability in neutrophils in blood or spleen (Fig. 3 and S1). These indicate that iGC is involved in phagocytosis and the deficiency of iGC impairs the defense of microbial infection during sepsis.

### Potential precision medicine in sepsis: GC supplementation improves the survival of septic mice with adrenal insufficiency but not septic mice with normal adrenal function

In this study, we showed that GC supplementation benefited septic mice with RAI (Figure. 4B), indicating that GC therapy might be potentially beneficial for a subgroup of septic patients having RAI. Moreover, GC supplementation did not benefit septic mice with normal adrenal function (Figure. 4A). This explains one of possibility why the efficacy and efficiency of GC therapy in septic patients are controversial. In our previous study, it also showed the same effect of GC supplementation in septic mice with or without RAI ^38^. Moreover, Ai et al. ^38^ indicated that early administration of GC improved more survival in septic mice with adrenal insufficiency. Recent clinical trials also showed that early GC therapy improves survival in septic patients. In this study, we administrated hydrocortisone in mice after CLP. The survival of septic mice with adrenal insufficiency increased in single dose of hydrocortisone treatment, showing that GC therapy benefits septic mice with adrenal insufficiency. However, the survival rate of septic mice with adrenal insufficiency treated with hydrocortisone decreased gradually with a time course. Considering that the duration of action of hydrocortisone is about 8-12 hours, this suggests that a single dose of hydrocortisone may not be enough to maintain the effect of GC on the protection against sepsis. Thus, the proper dose and number of dose should be investigated further. In addition, the method for the measurement of cortisol level and the criteria of adrenal insufficiency are still not standardized, which affects the accuracy of diagnosis of adrenal insufficiency and complicates the outcomes of sepsis treatment. These limitations impede and complicate clinical trials studying the efficacy of GC therapy in septic patients with adrenal insufficiency. Therefore, our adrenal insufficiency mouse model is pivotal for exploring more potential protective roles of GC therapy in precision medicine in sepsis treatment.

High concentration of IL-6 is associated with worse sepsis outcome such as early mortality ^56^. GC therapy enables to reduce the level of plasma pro-inflammatory cytokines in septic patients ^57–59^, showing the protective effects of GC therapy in sepsis. In this study, GC supplementation significantly reduced IL-6 production in mice with adrenal insufficiency at 4 and 20 h after CLP (Fig. 4C–4E). Although there was no significant difference in IL-6 level between two groups of SR-BI^fl/fl^ mice, GC supplementation increased IL-6 level in in SR-BI^fl/fl^ mice in sepsis (Fig. 4C-E). This implicated that the suppression of IL-6 during sepsis is one of the important protective effects of GC therapy to increase survival rate of septic mice with adrenal insufficiency. However, the increased plasma IL-6 level in septic mice with normal adrenal function which supplemented with hydrocortisone might aggravate the severity of sepsis in these mice.

Acute kidney injury is commonly seen in septic patients and associated with the increased rate of death ^60,61^. Ai et al. ^38^ showed the kidney injury in septic mice with adrenal insufficiency. So far, there are limited studies showing the roles of corticosteroids in sepsis-induced kidney injury. Johannes et. al ^62^ reported that low-dose dexamethasone-supplemented fluid resuscitation improved renal oxygen delivery and restored kidney function in Wistar rats treated with LPS, showing the potential application of corticosteroids in the prevention of sepsis-induced acute renal failure. Our data showed that the deficiency of iGC results in severe kidney injury and GC supplementation improves this injury during sepsis based on the plasma BUN (Fig. 4F and 4G). Moreover, GC treatment aggravates septic mice with normal adrenal function.

Taken together, GC therapy suppresses pro-inflammatory responses and ameliorates kidney injury, which improves the survival of septic mice with adrenal insufficiency. On the contrary, GC therapy aggravates the severity of septic mice with normal function might be because of the disruption of plasma corticosterone level (data not shown), increased the production of pro-inflammatory cytokines and exacerbating kidney injury. In addition to the limitations of diagnosis of adrenal insufficiency ^63^, the data implicate the possible reasons why the effects of GC therapy on survival of septic patients is not consistent ^25,64^.

### Stress level of GC (iGC) is required for the effective suppression of inflammatory cytokine production in sepsis

The higher concentration of GC (iGC) is stimulated and required for higher stress condition in sepsis (Fig. 1E and 2). Glucocorticoids play anti-inflammatory and immunosuppressive effects on macrophages. Our data showed that higher dose of hydrocortisone reduced more pro-inflammatory cytokines production; moreover, higher dose of hydrocortisone is required for the suppression of cytokine production that induced by higher concentration of LPS (Fig.5). IFN-γ is one of main macrophage activators ^48^ and IFN-γ-priming enhances the pro-inflammatory responses in macrophages. Higher dose of hydrocortisone is required for the suppression of cytokine production in LPS and IFN-γ stimulated macrophages (Fig.5). MKP-1 suppresses TLR-4-mediated inflammatory responses in macrophages through suppressing p38 MAPK ^49,50^. In addition to GC, IFN-γ inhibits MKP-1 expression in M-CSF-stimulated macrophages ^65,66^. However, these studies did not show the effects of GC on LPS and IFN-γ-stimulated macrophages. In our data, in the presence of IFN-γ, the effects of GC on mRNA expression of MKP-1 was compromised, implicating that the stress level of GC (iGC) is important for the suppression of activated macrophage in sepsis. These in vitro experiments support the important roles of stress level of GC (iGC) in the regulation of immune response in sepsis.

