## Supplementary material for "Relative adrenal insufficiency is a risk factor and an endotype of sepsis - A proof of concept study to support a precision medicine approach for glucocorticoid sepsis therapy": Supplemantl data

**Supplemental data**

**Table S1. List of cytokines production which did not show significant difference between SR-BI^fl/fl^ mice and SF1creSR-BI^fl/fl^ mice treated with CLP.** SR-BI^fl/fl^ mice and SF1creSR-BI^fl/fl^ mice were treated with CLP (25G, full ligation) for 4 and 20h. Plasma was harvested and analyzed for cytokines levels. Data represent means ± sem. n=6.

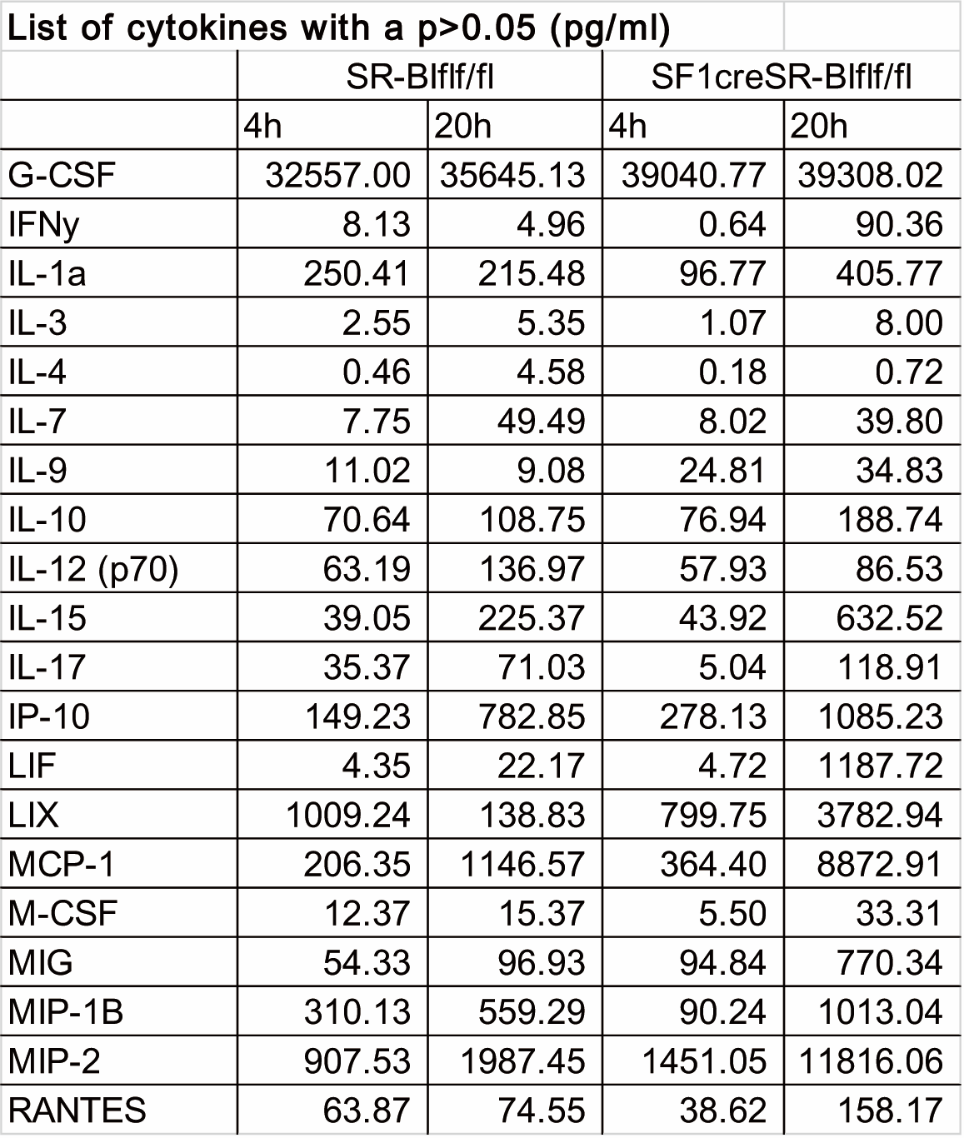

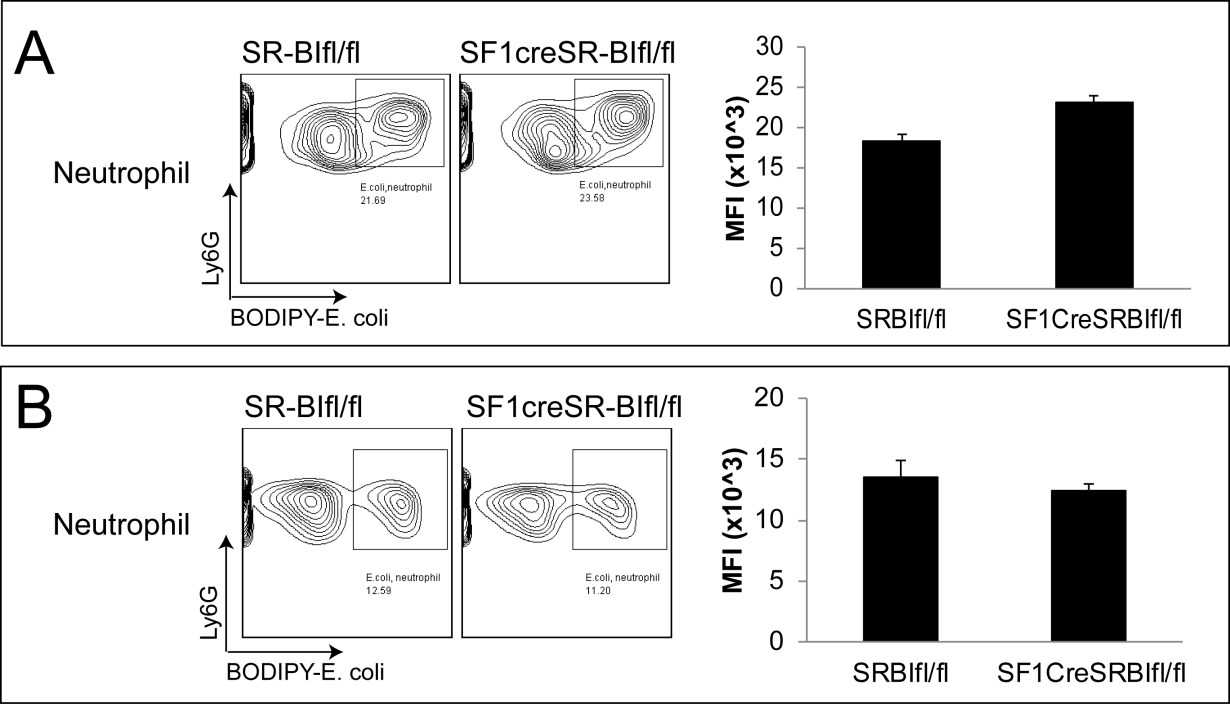

**FigS1. The effect of the deficiecny of iGC on the phagocytic ability of neutrophils in blood and spleen in sepsis.** SR-BI^fl/fl^ mice and SF1creSR-BI^fl/fl^ mice were treated with CLP for 17h and then injected with 10^9^ CFU/ml BODIPY-conjugated *E. coli* via i.v. for 1 hour. The MFI of *E. coli* in phagocytes in (A) spleen and (B) blood were analyzed with flow cytometry. n = 4–5. Data represent means ± sem.
